# Dopamine dysregulation in Parkinson’s disease flattens the pleasurable urge to move to musical rhythms

**DOI:** 10.1101/2023.02.27.530174

**Authors:** Victor Pando-Naude, Tomas Edward Matthews, Andreas Højlund, Sebastian Jakobsen, Karen Østergaard, Erik Johnsen, Eduardo A. Garza-Villarreal, Maria A. G. Witek, Virginia Penhune, Peter Vuust

**Affiliations:** Center for Music in the Brain, Department of Clinical Medicine, Aarhus University & The Royal Academy of Music Aarhus/Aalborg, Denmark; Center of Functionally Integrative Neuroscience, Department of Clinical Medicine, Aarhus University Hospital, Aarhus, Denmark; Department of Neurology, Aarhus University Hospital, Aarhus, Denmark; Instituto de Neurobiología, Universidad Nacional Autónoma de México (UNAM) campus Juriquilla, Queretaro, Mexico; Department of Music School of Languages, Cultures, Art History and Music, University of Birmingham, Birmingham, United Kingdom; Department of Psychology, Concordia University, 7141 Sherbrooke St W, Montreal, Quebec, Canada, H4B 1R6

**Keywords:** Parkinson’s Disease, music-induced pleasure, predictive processes, rhythm, harmony, dopamine, basal ganglia

## Abstract

The pleasurable urge to move to music (PLUMM) elicits activity in motor and reward areas of the brain and is thought to be driven by predictive processes. Dopamine within motor and limbic cortico-striatal networks is implicated in the predictive processes underlying beat-based timing and music-induced pleasure, respectively. This suggests a central role of cortico-striatal dopamine in PLUMM. This study tested this hypothesis by comparing PLUMM in Parkinson’s disease patients, healthy age-matched, and young controls. Participants listened to musical sequences with varying rhythmic and harmonic complexity (low, medium, high), and rated their experienced pleasure and urge to move to the rhythm. In line with previous results, healthy younger participants showed an inverted U-shaped relation between rhythmic complexity and ratings, with a preference for medium complexity rhythms, while age-matched controls showed a similar, but weaker, inverted U-shaped response. Conversely, PD patients showed a significantly flattened response for both the urge to move and pleasure. Crucially, this flattened response could not be attributed to differences in rhythm discrimination and did not reflect an overall decrease in ratings. Together, these results support the role of dopamine within cortico-striatal networks in the predictive processes that form the link between the perceptual processing of rhythmic patterns, and the affective and motor responses to rhythmic music.

## 1. Introduction

When listening to rhythmic music, we often feel an automatic and pleasurable urge to tap our feet or bob our heads along with the underlying beat or pulse of the rhythm. This pleasurable urge to move to music (PLUMM) is a key component of groove ^1–6^, and highlights the deep connection between music, movement, and pleasure. In a recent neuroimaging study, we showed that PLUMM is associated with greater activity in dorsal and ventral striatal regions within the basal ganglia (BG), along with cortical regions associated with motor and limbic cortico-striatal loops ^7,8^. This aligns with previous work implicating these loops in the predictive processes thought to drive both rhythm perception and music-induced pleasure, along with motor timing and reward processes more generally. For example, the motor cortico-striatal loop is consistently engaged during rhythm and beat perception, and production ^9– 14^, whereas the limbic cortico-striatal loop is implicated in music-induced pleasure ^15–21^. Within cortico-striatal networks, dopamine seems to have a functional role in both beat-based ^22–26^ and motor timing ^27^, as well as music-induced pleasure ^18,28,29^. This confluence of musically relevant timing, motor, and reward processes suggests a key role of cortico-striatal dopamine in PLUMM ^8^. To directly test this, we compared ratings of pleasure and wanting to move in two groups of Parkinson’s Disease (PD) patients, as well as age-matched and younger healthy controls.

Dopamine is thought to be necessary for the predictive processes that are fundamental to motor timing and reward computations ^30–32^, likely via separate pathways within motor and limbic cortico-striatal loops. The motor cortico-striatal loop, including the putamen, premotor cortex, and supplementary motor area, encompasses the nigrostriatal dopaminergic pathway, wherein dopaminergic cells in the substantia nigra pars compacta project to the dorsal striatum, including the putamen and caudate. Nigrostriatal dopamine is implicated in timing processes in the hundreds of milliseconds range that are necessary for sensory anticipation and motor preparation ^31,33–35^. The limbic cortico-striatal loop, including the nucleus accumbens and medial prefrontal cortices, encompasses the mesolimbic dopaminergic pathway, wherein cells in the ventral tegmental area project to the ventral striatum including the nucleus accumbens. Longstanding models suggest that mesolimbic dopamine codes for reward prediction errors, indicating whether a reward was better or worse than expected ^32^. Recent empirical and theoretical work suggest an expansion of the role of striatal dopamine to include sensory prediction errors (when sensory outcomes deviate from predictions), as well as the certainty of sensorimotor, state, or value representations ^36–38^.

Predictive processes are crucial to the perception and production of music, as well as affective responses including PLUMM. Music is often highly structured both in time and tonal space. Embellishments on, or deviations from this structure can elicit prediction confirmations or errors, and in turn, affective and motor responses within listeners ^39–44^. In terms of PLUMM, there is an inverted U-shaped relation between urge to move and pleasure ratings, and rhythmic complexity (predictability), wherein the highest ratings are elicited by rhythms that are neither too simple (predictable) nor too complex ^2,45–49^. A predictive processing account suggests that this pattern is driven by both the number of rhythm-based temporal prediction errors and the certainty of the antecedent predictions ^43,44,50,51^. According to this framework, predictions arise from an internal model based on prior experience. In the case of rhythmic music, the model is the meter, which, in western musical traditions is the pattern of strong beats (when a note is very likely to occur) and weak beats (when a note is less likely). Therefore, moderately complex, high-PLUMM rhythms are regular enough to enable relatively precise metrical predictions and complex enough to challenge them. Recently, harmonic complexity was also shown to influence PLUMM via its effect on pleasure, in contrast to rhythmic complexity which directly affects both the urge to move and pleasure ^46^. This suggests that the urge to move and pleasure can be at least partially disentangled via the differential effects of rhythm and harmony.

The role of dopamine in temporal and reward-related predictive processes is well established, however, it has yet to be linked to PLUMM. A recent theoretical model suggests that nigrostriatal dopamine plays a key role in probabilistic representations of the musical beat and thus supports beat-based motor and perceptual timing ^52^. This is consistent with studies involving patients with PD, which is characterized by cell loss in the substantia nigra leading to dysregulated dopamine in the nigrostriatal pathway and thus problems with motor-timing (e.g., dysfunctional gait and bradykinesia). Specifically, PD patients show poorer performance and altered brain activity during perceptual and motor tasks that rely on beat-based timing ^22–24,26,53–62^. More generally, PD patients show a reduced urge to engage with music via dancing or singing ^63^, suggesting a weakened link between music and movement.

The BG and dopamine are also implicated in music-induced pleasure ^18,28,29^. Activity in the limbic cortico-striatal loop, as well as functional and structural connectivity within this loop, and between the striatum and auditory cortex, are associated with music liking ^15–17^, music wanting (i.e., the money one is willing to spend on a given piece; Salimpoor et al., 2013), and music reward sensitivity ^64,65^. Activity in the nucleus accumbens has also been linked to the predictive processes thought to underlie music-derived pleasure ^19–21,66^. Evidence of a causal role for striatal dopamine in music-induced pleasure more generally comes from two parallel lines of research showing that modulations in BG activity or dopamine availability results in corresponding modulations in liking and wanting of music ^28,67^. However, many of these studies have focused on harmonic predictions rather than the temporal predictions that dominate in PLUMM (Matthews et al., 2019).

Given the role of dopamine within cortico-striatal networks in both rhythm-based predictive timing and music-derived pleasure, we investigated the link between cortico-striatal dopamine and PLUMM by comparing urge to move and pleasure ratings in PD patients along with healthy older and younger controls. We hypothesized that the inverted U-shaped relation between rhythmic complexity, wanting to move and pleasure would be altered in PD patients compared to controls but that the relationship between harmonic complexity and PLUMM would be unaffected. Aging has also been linked to changes in neural and behavioural correlates of rhythm ^68–74^ and reward processing ^75–78^. Therefore, the inclusion of both healthy older and younger controls also allowed us to examine any age-related changes in overall PLUMM response.

## 2. Material and Methods

### 2.1. Participants

The study was conducted at the Center for Music in the Brain, Aarhus University, Denmark. A total of 112 participants performed the experiment. Data from 101 of these participants were analysed after 11 PD patients were excluded from the analysis due to incomplete data. **Table 1** shows a summary of demographic data.

**Table 1.** Demographics and behavioural questionnaires.

|  | HC<sub>young</sub> | HC<sub>old</sub> | PD<sub>DA</sub> | PD<sub>nonDA</sub> |
| --- | --- | --- | --- | --- |
| Participants (n) | 27 | 26 | 23 | 24 |
| Gender (females) | 14 | 21 | 4 | 10 |
| Age (years) | 24.9±3.2 | 65.7±8.2 | 68.2±4.8 | 66.8±9 |
| Education (years) | 18.2±2.1*** | 14.7±2.3 | 14.9±2.9 | 14.9±3.5 |
| Musical Training (years) | 4.5±6.4** | 2.6±5.9 | 0.3±0.9 | 2±5.4 |
| Music Time (hr/week) | 22±14 | 19±15 | 18±18 | 16±15 |
| MDI | 3.1±3** | 5±6* | 8.4±4.6 | 8.1±5.7 |
| MoCA | 29±1.1 | 29±1.2 | 28±1.4 | 28±1.5 |
| PD diagnosis (years) | - | - | 6.6±4.7 | 5.6±4.9 |
| MDS-UPDRS-III | - | - | 29.7±11 | 25.2±12.3 |
| Hoehn and Yahr | - | - | 1.8±0.4 | 1.6±0.5 |
| MET: Melody | 36.7±5.8 | 34±5.6 | 31.5±4 | 32.4±5.4 |
| MET: Rhythm | 38±5.5 | 34±5.7 | 32.7±3.9 | 32.3±3.9 |

MET, musical ear test; MDI, major depression inventory; MoCA, Montreal cognitive assessment; MDS-UPDRS-III, motor part of the Movement Disorder Society Unified Parkinson's Disease Rating Scale. \*\*\* significantly different than all other groups, \*\* significantly different than both PD groups, \* sig different than PD<sub>DA</sub>.

PD patients (age 46-80) were diagnosed with idiopathic PD, in early- to mid-stages based on the Hoehn & Yahr scale ^79,80^, who were diagnosed by a certified neurologist according to the Movement Disorder Society (MDS) diagnostic criteria ^81^. PD patients whose symptoms exceeded stage III on the Hoehn & Yahr scale (Hoehn & Yahr, 1967) were excluded from the analysis to avoid confounding effects of cognitive impairments associated with late-stage PD. Additionally, patients were excluded if they were in treatment with deep brain stimulation or if they suffered from severe walking and balance problems, had important uncontrolled metabolic problems (e.g., diabetes, hypertension), hearing loss, or any other neurological or psychiatric disorders. Patients were instructed to maintain their regular medication regimen which included levodopa for all patients except for three who were only taking a dopamine agonist. Dopamine agonist medication is often taken in combination with levodopa and the combination can lead to disruptions in motivation and reward processes ^82^. To control for these effects, the PD patients were divided into two groups, those who were taking dopamine agonist medications (PD_DA_; n = 23) and those who were not (PD_nonDA_; n = 24). There were two groups of healthy controls; age-matched older adults (HC_old_; age 47-77; n = 27) and younger adults (HC_young_; age 21-33; n = 27). HCs had normal hearing and reported no metabolic, neurological, or psychiatric disorders. All participants in the HC_old_ group and in both PD groups reported their nationality as Danish. Nineteen participants in the HC_young_ group reported their nationality as Danish while the remaining participants reported their country of nationality as Poland, German, Egypt, Philippines, Netherlands, UK, Turkey, and Brazil.

All participants gave their consent verbally and in written form before the experiment. The study was conducted in accordance with the Declaration of Helsinki and approved by the Internal Review Board at the DNC (*Dansk Neuroforskningscenter*), Aarhus University Hospital, Aarhus, Denmark. Patients received no compensation for participating in the study, but the costs of transportation were covered. Healthy controls received 200 DKK, in compensation for their participation in the two-hour experiment.

### 2.2. Behavioural Questionnaires and Musical Ear Test

All participants filled out the Major depression inventory (MDI) to exclude depression ^83^ and the Montreal cognitive assessment (MoCA) to exclude dementia ^84^. PD patients filled out the MDS-Unified Parkinson’s disease rating scale (MDS-UPDRS-III) to ensure PD diagnosis ^85^. Authorization to use materials owned by the International Parkinson and Movement Disorder Society (MDS) was granted to author VPN, for research purposes only. Additionally, rhythm and melody discrimination ability was assessed in all participants via the Musical Ear Test (MET; ^86^. **Table 1** shows a summary of questionnaire and MET data.

### 2.3. PLUMM rating paradigm

Participants listened to and rated experienced pleasure and wanting to move for rhythmic stimuli drawn from previous studies in our lab ^8,46,49^. Stimuli varied in rhythmic complexity, measured via the syncopation index ^87,88^ which captures the degree to which a pattern deviates from the meter (in this case 4/4). There were three levels of rhythmic complexity (Low, Medium, and High) with three different rhythms in each level. The rhythmic patterns were made up of piano chords (six notes spanning four octaves in D Major) with a single chord repeating within pattern/sequence, along with an isochronous hihat (corresponding to eighth notes). The rhythms also varied across three levels of harmonic complexity, with three different chords per level. Harmonic complexity corresponded to the degree to which the chords deviated from the D Major key and was quantified as roughness using the MIR Toolbox in Matlab ^89–91^. The rhythms and chords were combined into six versions (i.e., six out of the nine possible combinations) of each rhythmic and harmonic complexity combination (e.g., high rhythmic complexity, low harmonic complexity), for a total of 54 stimuli. The sequences were presented at 96 bpm and lasted 10 seconds. All stimuli were created using Cubase (Steinberg Media Technologies).

At the start of the task, participants listened to two musical sequences that were not part of the experiment, to adjust the volume to a comfortable level. Participants listened to each sequence and rated pleasure and wanting to move on a 5-point Likert scale (1 = no pleasure/not wanting to move, 5 = very pleasurable/want to move), by using numbers on the computer keyboard to select their rating. The ratings were made according to the following questions: a) To what extent does this rhythm make you want to move? and b) How much pleasure do you experience listening to this rhythm? Following stimulus presentation, participants were given five seconds to make their rating. Ratings were made for all 54 stimuli in two consecutive sessions, wanting to move then pleasure, to avoid influences of one rating on the other. Within each session, each musical sequence was presented in a fully randomized order. Participants listened via Beyerdynamic DT 770 Pro headphones.

### 2.4. Statistical analysis

Only data from participants who completed all questionnaires, and all trials of the PLUMM rating paradigm were analysed. Therefore, the analyses described below are implemented on a complete data set with no missing values (N = 101).

#### Behavioural questionnaires and Musical Ear Test

Descriptive and inferential statistics of data were performed using R (version 4.1.1) in RStudio. Non-parametric (Kruskal-Wallis rank sum) tests were used to check for differences in demographic, cognitive, motor, and depression questionnaire responses between groups. Post-hoc analyses were conducted using the Dunn test of multiple comparisons corrected with the Bonferroni method.

The MET results showed a normal distribution; therefore, a one-way ANOVA was conducted to compare the groups. Post hoc analyses were conducted using the Tukey HSD multiple comparisons of means test with a 95% family-wise confidence level.

#### Wanting to move and pleasure ratings

Effects of group, rhythmic and harmonic complexity, and their interactions, on wanting to move and pleasure ratings were investigated using two separate regression models. Data from all four groups (HC_old_, HC_young_, PD_DA_, PD_nonDA_) were included in these models. As there were between-group differences in years of education, years of formal music training, and MDI score, these variables were included as covariates. The two models were submitted to separate Type III sum of squares analysis of variance (ANOVA) to test for main effects and interactions. If the ANOVA showed a significant or near-significant three-way interaction, F-tests were used to test for a rhythm by harmony interaction simultaneously in all groups. Second-order polynomial and pairwise contrasts were used to test quadratic and linear effects of rhythmic and harmonic complexity, and to compare these effects between groups, using the emmeans package (Lenth et al., 2018). For the ANOVAs, the degrees of freedom were estimated using the Satterthwaite method. For the follow-up contrasts, degrees of freedom were estimated using the Kenward-Roger method and the multivariate t method was used to correct for multiple comparisons.

Finally, we tested correlations between the ratings and PD motor symptom severity in PD_DA_ and PD_nonDA_, and age and MET scores in PD_DA_, PD_nonDA_, and HC_old_. A summary measure capturing the pattern of ratings (Medium – (Low + Medium)/2), calculated on estimated ratings from the models were used in the correlations. False discovery rate was used to correct for multiple correlations.

To account for inter-individual variability, as well as variability between versions of stimuli (items) within the same complexity levels, the regression models were estimated using linear mixed effects models on the raw pleasure and wanting to move ratings using lme4 (Bates et al., 2015) and afex (Singmann et al., 2013) packages. Random effects structures were determined following the recommendations of Bates et al. (2015) and Matuschek et al. ^92^. For both models, this resulted in by-participant random intercepts and random slopes for both rhythmic and harmonic complexity, as well as by-item random intercepts. Correlations between by-participant random slopes and intercepts were included for the analysis of wanting to move ratings but not pleasure ratings. Diagnostic plots of residuals indicated no violations of assumptions in either model.

## 3. Results

### 3.1. Demographic and Behavioural Questionnaires

Demographic and questionnaire data are presented in **Table 1**. A Kruskal-Wallis test indicated that PD_DA_, PD_nonDA_, and HC_old_ were matched in terms of age (*χ*^*2*^(2) = 1.21, *p* = .55) and MoCA scores (*χ*^*2*^(3) = 7.03, *p* = .07). Dunn tests showed no significant differences between PD_nonDA_ and PD_DA_ in years since PD diagnosis (*p* = .25), UPDRS (*p* = .23), or Hoehn & Yahr scale (*p* = .24). The groups differed in terms of gender ratio, with the HC_old_ being predominantly female and the PD groups being predominantly male. The groups also differed for years of education (*χ*^*2*^(3) = 25.66, *p* < .001), with HC_young_ having more education than all other groups (all *p* < .001). A significant effect of group on musical training (*χ*^*2*^(3) = 12.99, *p* = .004) was driven by HC_young_ having more years of musical training than both PD groups (both *p* < .05), but not HC_old_. There was no significant effect of group on hours spent interacting with music (*χ*^*2*^(3) = 3.98, *p* = .26). There was a significant effect of group on MDI scores (*χ*^*2*^(3) = 25.05, *p* < .001) driven by significantly lower scores in HC_old_ and HC_young_ compared to PD_DA_ (both *p* < .05) and HC_young_ compared to PD_nonDA_ (*p* = .001). As there were between-group differences in years of education, years of formal music training, and MDI score, these variables were included as covariates in the regression models.

A significant effect of group on the rhythm version of the MET (*χ*^*2*^(3) = 17.37, *p* < .001) was driven by significantly better performance in HC_young_ compared to all other groups (all *p* < .05). A significant effect of group on the melody version of the MET (*χ*^*2*^(3) = 12.48, *p* = .006) was driven by better performance in HC_young_ compared to PD_DA_ (*p* = .006) and PD_nonDA_ (*p* = .05), but not HC_old_ (*p* = .65).

### 3.2. Wanting to Move Ratings

#### Rhythmic complexity by group interaction

The ANOVA showed a significant group by rhythmic complexity interaction (**Table S1**) and no significant effects of the control variables (years of education, years of musical training and MDI). Follow up tests showed a significant quadratic effect in both HC_young_ and HC_old_, with HC_young_ showing a stronger effect compared to HC_old_ (*b*(97) = -1.216, 95% CI [-1.840, -0.589]; **Table S2** and **Figure 1A**). The quadratic effect was significantly stronger in HC_old_ compared to both PD_nonDA_ (*b*(97)= -1.066, 95% CI [-1.821, -0.312]) and PD_DA_ (*b*(97) = -1.113, 95% CI [-1.876, -0.349]). PD_nonDA_ showed a significant negative linear effect which was driven by significantly higher ratings for low compared to high complexity rhythms (*b*(85.3) = 0.468, 95% CI [0.091, 0.845]). PD_DA_ showed a relatively flat pattern of ratings as reflected in no significant linear or quadratic effects. The difference in linear effects between PD_nonDA_ and PD_DA_ was not significant.

**Figure 1.**
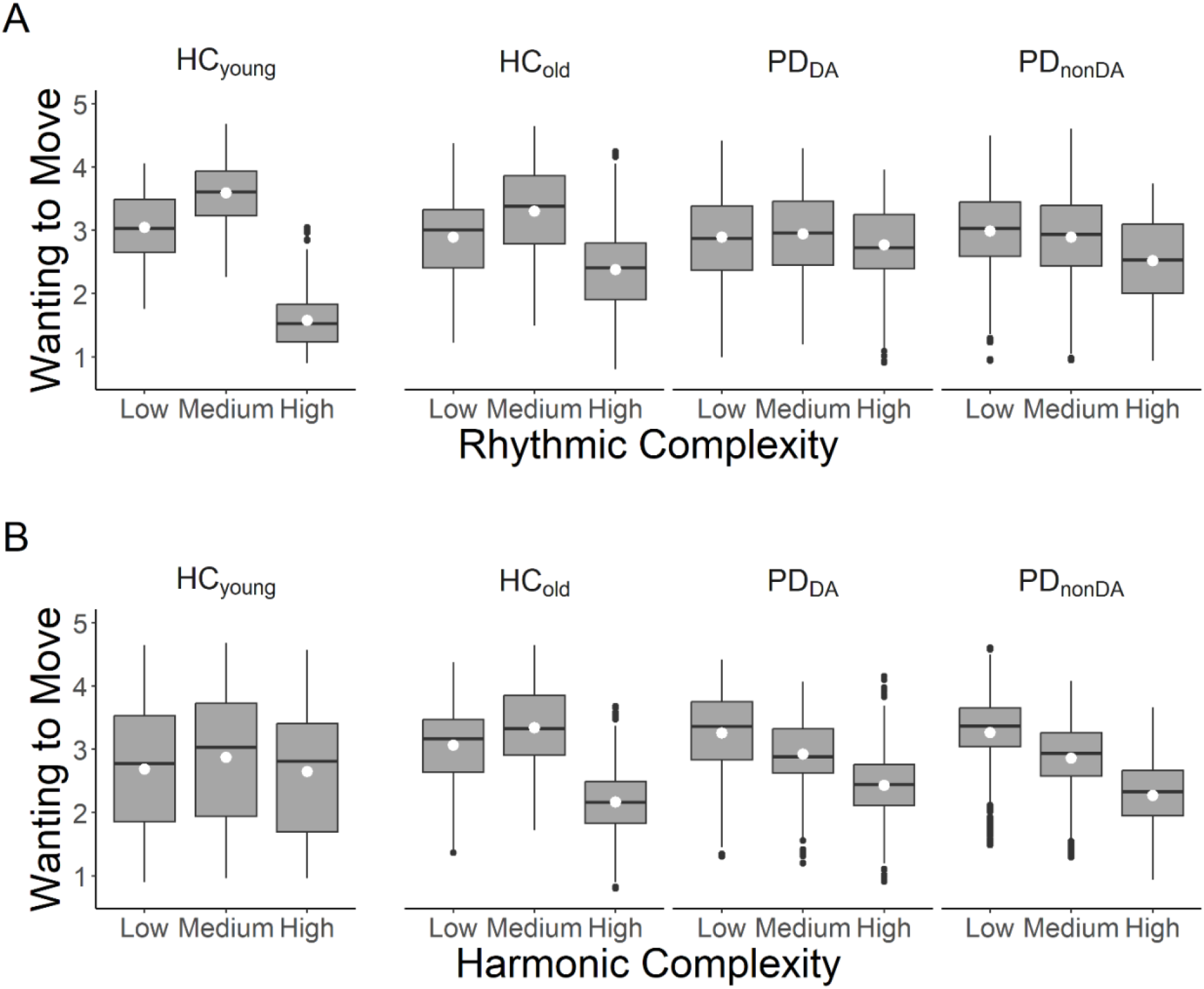
The effects of rhythmic (A) and harmonic complexity (B) on Wanting to Move ratings across groups. Values are estimated ratings from regression model. Center line, median; box limits, upper and lower quartiles; whiskers, 1.5x interquartile range; white points, means; black points, outliers.

#### Harmonic complexity by group interaction

The ANOVA also showed a group by harmonic complexity interaction (**Table S3**). Follow up tests showed that both HC_old_ and HC_young_ and that this effect was stronger in HC_old_ compared to HC_young_ (*b*(97) = 1.043, 95% CI [0.593, 1.490]; **Table S3** and **Figure 1B**). HC_old_ also showed a stronger quadratic effect compared to both PD_nonDA_ (*b*(97) = -1.261, 95% CI [-1.801, -0.720]) and PD_DA_ (*b*(97) = -1.289, 95% CI [-1.836, - 0.742]). As can be seen in **Figure 1B**, both PD_nonDA_ and PD_DA_ showed significant negative linear effects. This was reflected by significant low – medium (PD_nonDA_: *b*(89.2) = 0.403, 95% CI [0.153, 0.653]; PD_DA_: (*b*(90.1) = 0.333, 95% CI [0.078, 0.589]) and medium – high (PD_nonDA_: *b*(89.0) = 0.593, 95% CI [0.343, 0.842]; PD_DA_: (*b*(90.0) = 0.495, 95% CI [0.241, 0.749]) contrasts. There were no significant differences in these effects between PD_nonDA_ and PD_DA_.

#### Rhythmic complexity by harmonic complexity interaction

The ANOVA also showed a near-significant three-way interaction (**Table S1**). F-tests revealed a significant rhythmic complexity by harmonic complexity interaction in HC_old_ (*F*(4, 4908.05) = 4.38, *p* = .002) and a near-significant interaction (*F*(4, 4908.05) = 2.11, *p* = .077) for HC_old_, but not for PD_DA_ or PD_nonDA_ (both *p* > .2). Follow up contrasts in HC_old_ showed that the quadratic effect of rhythmic complexity was significantly stronger, here reflecting a more peaked inverted U, for medium complexity chords compared to both low (*b*(4908) = -0.611, 95% CI [-1.177, -0.045]) and high (*b*(4908) = -0.833, 95% CI [-1.400, -0.267]) complexity chords. Unlike HC_old_, HC_young_ showed no significant differences in the quadratic effect of rhythmic complexity between medium and low or medium and high complexity chords.

#### Main effect of group

The ANOVA showed that the main effect of group was not significant (*F*(3, 94.1) = 0.78, *p* = 0.508). Therefore, although the groups showed different patterns in their ratings, there was no overall difference when averaging over rhythmic and harmonic complexity.

#### Correlations between Wanting not Move ratings, UPDRS, and MET scores

In PD patients, correlations between UPDRS and the contrast estimates for both rhythmic (*r*(47) = -0.24, 95% CI [-0.52, 0.09]) and harmonic complexity were not significant (*r*(47) = -0.24, 95% CI [-0.52, 0.09]). Correlations with MET scores were also not significant (all *p* > .09).

### 3.3. Pleasure ratings

#### Rhythmic complexity by group interaction

The ANOVA showed significant group by rhythmic complexity interaction (**Table S4**) and no significant effects of the control variables. Follow up tests showed a stronger quadratic effect in HC_young_ compared to HC_old_ (*b*(98.5) = -1.128, 95% CI [-1.87, -0.389]) and no significant quadratic effects in HC_old_, PD_nonDA_ and PD_DA_ (**Table S5** and **Figure 2A**). PD_nonDA_ showed a significant negative linear effect (*b*(119.6) = -0.602, 95% CI [-1.077, -0.127]) driven by higher ratings in medium compared to high complexity rhythms (*b*(117.5) = 0.565, 95% CI [0.091, 1.039]). The same contrast was also significant in HC_old_ (*b*(113.3) = 0.512, 95% CI [0.060, 0.964]). There were no significant differences in the linear effect between groups. As can be seen in **Figure 2A**, both HC_old_ and PD_nonDA_ show similar ratings for low and medium complexity rhythms, then a drop off for high complexity rhythms. Conversely, PD_DA_’s ratings are relatively flat across all levels.

**Figure 2.**
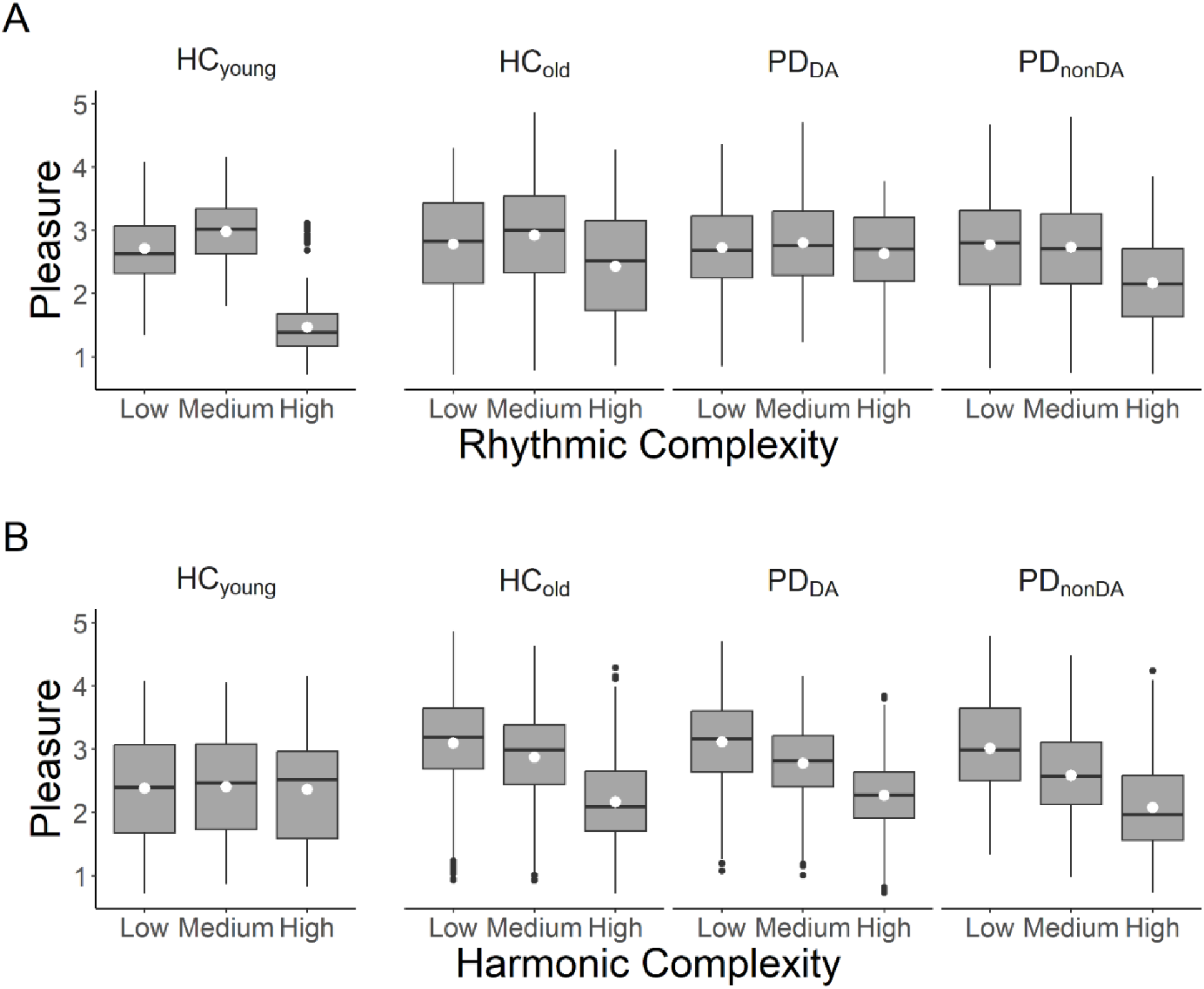
The effects of rhythmic (A) and harmonic complexity (B) on Pleasure ratings across groups. Values are estimated ratings from regression model. Center line, median; box limits, upper and lower quartiles; whiskers, 1.5x interquartile range; white points, means; black points, outliers.

#### Harmonic complexity by group interaction

The ANOVA also showed a significant group by harmonic complexity interaction (**Table S4**). Follow up tests indicated that HC_old_ showed greater linear (*b*(103) = -0.93, 95% CI [-1.320, -0.541]) and quadratic (*b*(103) = -0.44, 95% CI [-0.850, -0.030]) effects compared to HC_young_, in whom neither effects were significant. As can be seen in **Figure 2B**, HC_old_, PD_nonDA_ and PD_DA_ all show a negative linear pattern as supported by a significant linear contrast in these groups (**Table S6**). These three groups showed significant differences between low and medium and medium and high complexity chords. However, HC_old_ showed a smaller difference between low and medium chords leading to a significant quadratic effect. Follow up test showed a significant quadratic effect in HC_old_ and significant negative linear effects for both PD_nonDA_ and PD_DA_. There were no significant differences between groups in either the quadratic or linear contrasts.

#### Rhythmic complexity by harmonic complexity interaction

The ANOVA also showed a significant three-way interaction (**Table S4**). Follow up F-tests showed that PD_nonDA_ showed a near-significant rhythmic complexity by harmonic complexity interaction (*F*(4, 4914.92) = 2.245, *p* = 0.061). PD_DA_ and HC_old_ did not show a significant or near-significant interaction (both p > .16). Follow up contrasts in PD_nonDA_ showed higher ratings for medium complexity rhythms compared to high complexity rhythms for both low (*b*(159) = 0.660, 95% CI [0.136, 1.183]) and medium (*b*(159) = 0.646, 95% CI [0.123, 1.168]), but not high, complexity chords. There was no significant difference between medium and low complexity rhythms at any level of harmonic complexity.

Comparing HC_young_ and HC_old_, F-tests showed that this interaction was significant in HC_young_ (*F*(4, 4914.92) = 4.464, *p* =.001) but not in HC_old_ (*F*(4, 4914.92) = 0.422, *p* = 0.793). Follow up contrasts showed that the quadratic effect of rhythm in HC_young_ was stronger for low compared to high complexity chords (*b*(4915) = -0.735, 95% CI [-1.300, -0.170]), but no differences in the quadratic effect of rhythm between low vs medium and medium vs high complexity chords.

#### Main effect of group

The ANOVA also showed a main effect of group (**Table S4**). There were no significant differences between PD_nonDA_, PD_DA_ and HC_old_. However, HC_old_ showed higher ratings overall compared to HC_young_ (*b*(94) = 0.453, 95% CI [0.141, 0.765]).

#### Correlations between Pleasure ratings, UPDRS, and MET scores

In PD patients, there was a significant negative correlation between UPDRS and rhythmic complexity contrast estimates (*r*(47) = -0.37, 95% CI [-0.62, -0.05]; **Figure 3**) but not harmonic complexity contrast estimates (*r*(47) = -0.03, 95% CI [-0.35, 0.30]). This indicates that as PD symptoms worsen the quadratic effect of rhythmic complexity flattens out and/or becomes more linear. No other correlations were significant.

**Figure 3.**
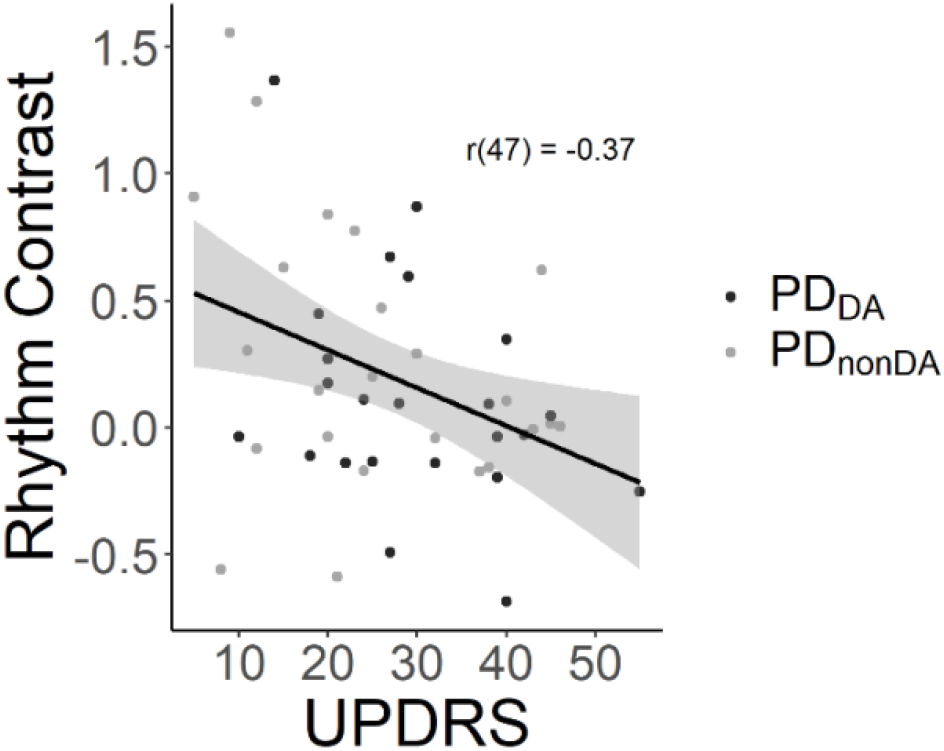
Correlation between contrast estimates (medium – (low + high)/2) from the effect of rhythmic complexity on pleasure, and severity of PD-related motor symptoms (UPDRS).

## 4. Discussion

Our results support a crucial role of cortico-striatal circuits and dopamine in the pleasurable urge to move to music ^8,93^. Both PD groups showed a flattening of the U-shaped relation between rhythmic complexity and ratings of pleasure and wanting to move compared to healthy age-matched controls. In addition, the severity of motor symptoms in PD was associated with a flatter effect of rhythmic complexity on pleasure. The inverted U-shaped relation was significantly weaker in healthy older compared to younger adults, suggesting that aging also influences the effect of rhythmic complexity on PLUMM. In addition, both PD groups showed a strong negative linear effect of harmonic complexity for both pleasure and wanting to move in comparison to healthy older adults. Importantly, PD patients did not differ from age-matched controls on a test of rhythm discrimination and did not show overall lower ratings for wanting to move and pleasure but rather a difference in the pattern of ratings. These results indicate that the flattening of the PLUMM response cannot easily be attributed to deficits in rhythm perception or a general inability to experience PLUMM. Finally, PD patients taking a dopamine agonist in addition to the standard levodopa regime generally showed a similar flattening effect to the PD_nonDA_ group. Taken together, these findings indicate that the chronic dysregulation of cortico-striatal dopamine associated with PD flattens the typical U-shaped relationship between rhythmic complexity and PLUMM. We propose that this points to the crucial involvement of cortico-striatal dopamine in the beat- and meter-based predictive processes thought to drive PLUMM.

### Rhythmic complexity

The flattened response to rhythmic complexity in PD patients supports a role of cortico-striatal dopamine in the effects of rhythm on both the urge to move and pleasure. The motor cortico-striatal loop is part of a core timing network, along with prefrontal and cerebellar regions ^94–96^, and is implicated in in relative or beat-based timing in particular ^9–14,56,97–102^. This is consistent with a role of the motor system in driving temporal predictions in the context of auditory perception and attention more generally ^103–107^. Involvement of the motor cortico-striatal loop implies a role of nigrostriatal dopamine. However, studies on beat perception in PD patients show mixed results ^22–24,26,53,54^. Conversely, many of these same studies show deficits in rhythm discrimination in PD patients ^22–24,26,53,54^, which could suggest that the flatter pattern of ratings seen here may be due to reduced ability to distinguish the stimuli. However, HC_old_, PD_nonDA_, and PD_DA_ significantly differed in their patterns of ratings but not in their rhythm discrimination ability, as measured by the MET. Further, there was no significant associations between MET scores and the patterns of ratings, or between MET scores and the severity of PD-related motor symptoms (UPDRS). Conversely, higher UPDRS was associated with a flatter effect of rhythmic complexity on pleasure. Therefore, our results suggest that PD disrupts the beat and meter-based predictive processes that form the link between perceptual processing of rhythmic patterns and the affective responses to rhythmic music.

According to predictive processing treatments of PLUMM, strongly weighted prediction errors, in the form of syncopations, drive the inverted U-shaped relation between rhythmic complexity and both the urge to move and pleasure ^43,44,50,51^. In this context, strongly weighted syncopations drive the urge to move, either directly to reinforce the metrical model, or indirectly via covert adjustments to that model. In terms of pleasure, by challenging our metric models, these syncopations potentiate model improvement ^93^, which is itself intrinsically rewarding ^108,109^. The weight of prediction errors due to syncopations depends on both the rhythmic context ^110^ and the strength of the listener’s metrical model ^111^. According to a recent theoretical model (Cannon & Patel, 2020), nigrostriatal dopamine underlies both the timing and certainty of beat-based predictions. Therefore, one interpretation of our results is that disrupted cortico-striatal dopamine in PD results in a weaker metrical model and more uncertain predictions, reducing pleasure and the urge to move for moderately complex rhythms. Similarly, high complexity rhythms are less disruptive to an already weak model and are thus less aversive, leading to a flatter pattern of ratings overall. Support for the role of metric model strength in PLUMM comes from studies comparing musicians and non-musicians. Behavioural and neural evidence suggests that musicians show stronger and/or more refined metrical models compared to non-musicians ^111–117^, and that musicians have a sharper U-shaped pattern of PLUMM ratings and greater preference for higher levels of complexity (Matthews et al., 2019; 2022). In the current study, healthy aging was also linked to a flatter effect of rhythmic complexity, particularly for pleasure ratings. Like PD, healthy aging is associated with differences in the neural correlates of predictive rhythmic pattern perception ^68^, an effect that is also linked to dopamine ^70^. Therefore, metrical model strength, supported by dopaminergic signalling, may provide a common mechanism underlying the effects of PD, aging, and musicianship on the shape and location of the inverted U relation between PLUMM and rhythmic complexity.

Neuroimaging evidence suggests that neural oscillations in the beta range (13-30 Hz) may encode the metrical model within the motor cortico-striatal loop and thus may provide a potential neural mechanism underlying the altered link between rhythm perception and affective responses in PD. Beta dysfunction within the BG is a hallmark of PD ^118^ suggesting a functional link between dopamine and beta activity. One proposal is that, by encoding the certainty (or salience) of state, sensorimotor, or value representations (e.g., rhythmic context; Friston et al, 2014), dopamine signals the need for motor preparation or resource allocation which are governed by beta modulations ^119^. Crucially, cortico-striatal beta activity is linked to beat and meter representations ^120–124^ and is sensitive to musical training ^125,126^. Further, PD patients show weaker beta entrainment in motor regions during rhythmic auditory and multimodal perceptual tasks ^127,128^, an effect that is reversed following treatment with levodopa and deep brain stimulation ^129^. Finally, the degree of beta dysfunction in cortico-striatal networks is correlated with the severity of motor symptoms in PD ^130^ which aligns with the association between severity of motor symptoms and the flattening of the effect of rhythmic complexity on pleasure ratings seen here. Together, these results point to a functional link between dopamine, beta activity, metrical model strength, and PLUMM, however, further work is necessary to solidify and clarify this link.

An alternative interpretation for the flattened effect of rhythmic complexity on PLUMM ratings in PD is that, rather than weakening the metrical model, dopamine dysregulation disrupts the signalling of syncopation-related prediction errors, thus reducing the urge to move and pleasure thought to result from these prediction errors. However, (reward) prediction errors are associated with mesolimbic dopamine ^32^ while PD primarily effects nigrostriatal dopamine, at least in early and mid-stages of the disease ^131^. One challenge is that the effects on metrical model strength would also be expected to affect the prediction errors, as the certainty of the meter-based prediction is thought to directly relate to the weight of the prediction error. Using tasks such as those from Palmer & Krumhansl (1990) or Perna et al. (2018), future work could explicitly test whether metrical representations do in fact differ between PD patients and healthy controls matched in age and musical training.

Interestingly, both PD groups showed a very similar pattern of ratings, aside from a drop in the urge to move and pleasure ratings in PD_nonDA_ for high complexity rhythms. This constitutes an internal replication and suggests that dopamine agonist medication does not have a strong effect on ratings over-and-above the general DA dysregulation and counteracting effects of levodopa present for all patients. To further delineate the role of dopamine, future work should compare ratings in both PD patients and healthy controls on and off levodopa medication.

### Harmonic Complexity

Contrary to our hypothesis, and in contrast to the pattern of ratings in young healthy adults shown here and elsewhere ^8,46^, PD patients and healthy older adults showed a greater preference for low complexity chords. A similar pattern was seen in a recent study wherein non-musicians preferred low complexity chords to medium, high, or octave chords, while musicians preferred medium complexity chords ^132^. Therefore, as with rhythmic complexity, PD and (lack of) musical training seem to show similar effects on chord preferences, again, possibly driven by predictive processes, albeit at a different temporal scale to metric predictions as the chords were static within each trial. Another possibility is that PD patients have an altered association between the complexity of a single chord and the emotion evoked by or referenced by this chord. For example, PD patients show differences in the ability to recognize emotions expressed in music ^133^, which is relevant as chords of varying complexity are often used to convey different emotions. However, the specific nature of this effect is so far inconclusive, as some studies show deficits in recognizing happiness and peacefulness in PD patients ^134^ and others show deficits in recognizing fear and anger ^135^. Other factors including listening habits and genre preferences may also have influenced the pattern of ratings seen here, however, research into these effects is lacking.

## 5. Conclusion

Overall, our results are consistent with models implicating the cortico-striatal circuits in PLUMM ^8,93^ and provide insight into the function of dopamine within these circuits more generally. We propose that dopamine depletion within the motor cortico-striatal loop disrupts beat- and meter-based predictive processes that are supported by beta modulations and are crucial for the affective response to rhythmic music. Beyond music, this proposal aligns with and complements theoretical and empirical work suggesting that premotor beta modulations transmit top-down predictive signals that temporally structure auditory perception, e.g., in the context of speech ^103,105,106,136,137^. Importantly, the current results better our understanding of dopamine’s role in linking temporal predictions and reward processing which can be leveraged to improve treatments for PD and other dopamine-related disorders.

## Author contributions

Victor Pando-Naude: Conceptualization, Methodology, Investigation, Formal analysis, Writing – Original Draft, Writing – Review & Editing, Supervision, Project Administration

Tomas Edward Matthews: Conceptualization, Formal analysis, Writing – Original Draft, Writing – Review & Editing, Visualization

Andreas Højlund: Conceptualization, Methodology

Sebastian Jakobsen: Investigation, Recruitment, Data acquisition

Karen Østergaard: Conceptualization

Erik Jonhnsen: Investigation, Supervision

Eduardo A. Garza-Villarreal: Conceptualization, Supervision

Maria Witek: Conceptualization, Supervision

Virginia Penhune: Conceptualization, Supervision, Writing – Review & Editing

Peter Vuust: Conceptualization, Supervision, Writing – Review & Editing, Supervision, Funding acquisition

## Acknowledgements

The Center for Music in the Brain (MIB) is supported by the Danish National Research Foundation (grant number DNRF 117). The authors would like to thank the students who assisted in the acquisition of data. Victor Poulsen, Anders Weile Larsen, Caroline Steen Sundberg, and Agata Patyczek. Special thanks to all patients and healthy controls for their participation in this study.

## Data availability

The data supporting the findings of this study is freely available at the Open Science Framework (OSF) website: https://osf.io/fszqe/?view_only=cffadc85e4004c8789ca5cf604f172c7

## Conflict of interest

The authors declare no conflict of interest for this work.

## Notes

https://osf.io/fszqe/?view_only=cffadc85e4004c8789ca5cf604f172c7

### Competing Interest Statement

The authors have declared no competing interest.

### Summary of Updates

Typo in abstract

## References

1. Janata, P., Tomic, S. T. & Haberman, J. M. Sensorimotor coupling in music and the psychology of the groove. J. Exp. Psychol. Gen. 141, 54–75 (2012).

2. Witek, M. A. G., Clarke, E. F., Wallentin, M., Kringelbach, M. L. & Vuust, P. Syncopation, body-movement and pleasure in groove music. PloS One 9, 1–12 (2014).

3. Duman, D., Snape, N., Toiviainen, P. & Luck, G. Redefining Groove. in Conference on Interdisciplinary Musicology (2022).

4. Câmara, G. S. & Danielsen, A. “Groove,” in The Oxford Handbook of Critical Concepts in Music Theory, eds A. Rehding and S. Rings (Oxford: Oxford University Press), 271–294. Cannon,. in The Oxford Handbook of Critical Concepts in Music Theory (eds. Rehding, A. & Rings, S.) 271–294 (2018). doi:10.1093/oxfordhb/9780190454746.013.17.

5. Madison, G. Experiencing Groove Induced by Music: Consistency and Phenomenology. Music Percept. 24, 201–208 (2006).

6. Senn, O. et al. Preliminaries to a psychological model of musical groove. Front. Psychol. 10, 1228 (2019).

7. Alexander, G., DeLong, M. R. & Strick, P. L. Parallel Organization of Functionally Segregated Circuits Linking Basal Ganglia and Cortex. Annu. Rev. Neurosci. 9, 357–381 (1986).

8. Matthews, T. E., Witek, M. A. G., Lund, T., Vuust, P. & Penhune, V. B. The sensation of groove engages motor and reward networks. NeuroImage 214, 1–12 (2020).

9. Bengtsson, S. L. et al. Listening to rhythms activates motor and premotor cortices. Cortex 45, 62–71 (2009).

10. Chen, J. L., Penhune, V. B. & Zatorre, R. J. Listening to musical rhythms recruits motor regions of the brain. Cereb. Cortex 18, 2844–2854 (2008).

11. Grahn, J. A. & Rowe, J. B. Feeling the beat: premotor and striatal interactions in musicians and nonmusicians during beat perception. J. Neurosci. 29, 7540–7548 (2009).

12. Grahn, J. A. & Rowe, J. B. Finding and feeling the musical beat: striatal dissociations between detection and prediction of regularity. Cereb. Cortex N. Y. N 1991 23, 913–21 (2013).

13. Schubotz, R. I., Friederici, A. D. & von Cramon, D. Y. Time perception and motor timing: a common cortical and subcortical basis revealed by fMRI. NeuroImage 11, 1–12 (2000).

14. Kasdan, A. V. et al. Identifying a brain network for musical rhythm: A functional neuroimaging meta-analysis and systematic review. Neurosci. Biobehav. Rev. 136, 104588 (2022).

15. Blood, A. J., Zatorre, R. J., Bermudez, P. & Evans, A. C. Emotional responses to pleasant and unpleasant music correlate with activity in paralimbic brain regions. Nat. Neurosci. 2, 382–387 (1999).

16. Blood, A. J. & Zatorre, R. J. Intensely pleasurable responses to music correlate with activity in brain regions implicated in reward and emotion. Proc. Natl. Acad. Sci. U. S. A. 98, 11818–11823 (2001).

17. Salimpoor, V. N. et al. Interactions between the nucleus accumbens and auditory cortices predict music reward value. Science 340, 216–9 (2013).

18. Salimpoor, V. N., Benovoy, M., Larcher, K., Dagher, A. & Zatorre, R. J. Anatomically distinct dopamine release during anticipation and experience of peak emotion to music. Nat. Neurosci. 14, 257–62 (2011).

19. Cheung, V. K. M. et al. Uncertainty and Surprise Jointly Predict Musical Pleasure and Amygdala, Hippocampus, and Auditory Cortex Activity. Curr. Biol. 1–9 (2019) doi:10.1016/j.cub.2019.09.067.

20. Gold, B. P., Mas-Herrero, E., Dagher, A. & Zatorre, R. J. Toward a fuller understanding of reward prediction errors and their role in musical pleasure. Proc. Natl. Acad. Sci. U. S. A. 116, 20815–20816 (2019).

21. Shany, O. et al. Surprise-related activation in the nucleus accumbens interacts with music-induced pleasantness. Soc. Cogn. Affect. Neurosci. 1–12 (2019) doi:10.1093/scan/nsz019.

22. Grahn, J. A. & Brett, M. Impairment of beat-based rhythm discrimination in Parkinson’s disease. Cortex 45, 54–61 (2009).

23. Cameron, D. J., Pickett, K. A., Earhart, G. M. & Grahn, J. A. The Effect of Dopaminergic Medication on Beat-Based Auditory Timing in Parkinson’s Disease. Front. Neurol. 7, 1–8 (2016).

24. Hsu, P., Ready, E. A. & Grahn, J. A. The effects of Parkinson’s disease, music training, and dance training on beat perception and production abilities. PLoS ONE 17, 1–16 (2022).

25. Geiser, E. & Kaelin-Lang, A. The function of dopaminergic neural signal transmission in auditory pulse perception: Evidence from dopaminergic treatment in Parkinson’s patients. Behav. Brain Res. 225, 270–275 (2011).

26. Biswas, A., Hegde, S., Jhunjhunwala, K. & Pal, P. K. Two sides of the same coin: Impairment in perception of temporal components of rhythm and cognitive functions in Parkinson’s disease. Basal Ganglia 6, 63–70 (2016).

27. Jahanshahi, M. et al. Dopaminergic modulation of striato-frontal connectivity during motor timing in Parkinson’s disease. Brain 133, 727–745 (2010).

28. Ferreri, L. et al. Dopamine modulates the reward experiences elicited by music. PNAS 116 (9) 37, 3793–3798 (2019).

29. Gebauer, L., Kringelbach, M. L. & Vuust, P. Ever-changing cycles of musical pleasure: The role of dopamine and anticipation. Psychomusicology Music Mind Brain 22, 152–167 (2012).

30. Tomassini, A., Ruge, D., Galea, J. M., Penny, W. & Bestmann, S. The Role of Dopamine in Temporal Uncertainty. J. Cogn. Neurosci. 28, 96–110 (2016).

31. Meck, W. H. Neuroanatomical localization of an internal clock: A functional link between mesolimbic, nigrostriatal, and mesocortical dopaminergic systems. Brain Res. 1109, 93–107 (2006).

32. Schultz, W., Dayan, P. & Montague, P. R. A neural substrate of prediction and reward. Science 275, 1593–1599 (1997).

33. Coull, J. T., Hwang, H. J., Leyton, M. & Dagher, A. Dopamine Precursor Depletion Impairs Timing in Healthy Volunteers by Attenuating Activity in Putamen and Supplementary Motor Area. J. Neurosci. 32, 16704–16715 (2012).

34. Jones, C. R. G. & Jahanshahi, M. Contributions of the Basal Ganglia to Temporal Processing: Evidence from Parkinson’s Disease. Timing Time Percept. 2, 87–127 (2014).

35. Behroozmand, R. & Johari, K. Sensorimotor Impairment of Speech and Hand Movement Timing Processing in Parkinson’s Disease. J. Mot. Behav. 51, 561–571 (2019).

36. Gershman, S. J. & Uchida, N. Believing in dopamine. Nat. Rev. Neurosci. 20, 703–714 (2019).

37. Friston, K. J. et al. The anatomy of choice: Dopamine and decision-making. Philos. Trans. R. Soc. B Biol. Sci. 369, (2014).

38. Friston, K. J. et al. Dopamine, affordance and active inference. PLoS Comput. Biol. 8, e1002327 (2012).

39. Huron, D. Sweet Anticipation: Music and the Psychology of Expectation. MIT Press (2006). doi:10.1525/mp.2007.24.5.511.

40. Meyer, L. B. Emotion and Meaning in Music. (University of Chicago Press, 1956).

41. Salimpoor, V. N., Zald, D. H., Zatorre, R. J., Dagher, A. & McIntosh, A. R. Predictions and the brain: how musical sounds become rewarding. Trends Cogn. Sci. 19, 86–91 (2015).

42. Zald, D. H. & Zatorre, R. J. On music and reward. in The neurobiology of Sensation and Reward (ed. Gottfried, J.) (Taylor & Francis, 2011).

43. Koelsch, S., Vuust, P. & Friston, K. Predictive Processes and the Peculiar Case of Music. Trends Cogn. Sci. 23, 63–77 (2019).

44. Vuust, P., Heggli, O. A., Friston, K. J. & Kringelbach, M. L. Music in the brain. Nat. Rev. Neurosci. 23, 287–305 (2022).

45. Sioros, G., Miron, M., Davies, M., Gouyon, F. & Madison, G. Syncopation creates the sensation of groove in synthesized music examples. Front. Psychol. 5, 1–10 (2014).

46. Matthews, T. E., Witek, M. A. G., Heggli, O. A., Penhune, V. B. & Vuust, P. The sensation of groove is affected by the interaction of rhythmic and harmonic complexity. PLoS ONE 14, 1–17 (2019).

47. Matthews, T. E., Witek, M. A. G., Thibodeau, J. L. N., Vuust, P. & Penhune, V. B. Perceived motor synchrony with the beat is more strongly related to groove than measured synchrony. Music Percept. 39, 423–442 (2022).

48. Spiech, C., Sioros, G., Endestad, T., Danielsen, A. & Laeng, B. Pupil drift rate indexes groove ratings. Sci. Rep. 12, 11620 (2022).

49. Stupacher, J., Wrede, M. & Vuust, P. A brief and efficient stimulus set to create the inverted U-shaped relationship between rhythmic complexity and the sensation of groove. PloS One 17, e0266902 (2022).

50. Vuust, P., Witek, M., Dietz, M. & Kringelbach, M. L. Now You Hear It: A predictive coding model for understanding rhythmic incongruity. Ann. N. Y. Acad. Sci. 1423, 19–29 (2018).

51. Vuust, P. & Witek, M. A. G. Rhythmic complexity and predictive coding: A novel approach to modeling rhythm and meter perception in music. Front. Psychol. 5, 1–14 (2014).

52. Cannon, J. J. & Patel, A. D. How Beat Perception Co-opts Motor Neurophysiology. Trends Cogn. Sci. 25, 137–150 (2020).

53. Benoit, C.-E. et al. Musically Cued Gait-Training Improves Both Perceptual and Motor Timing in Parkinson’s Disease. Front. Hum. Neurosci. 8, 1–11 (2014).

54. Geiser, E. & Kaelin-Lang, A. The function of dopaminergic neural signal transmission in auditory pulse perception: Evidence from dopaminergic treatment in Parkinson’s patients. Behav. Brain Res. 225, 270–275 (2011).

55. Bellinger, D., Altenmüller, E. & Volkmann, J. Perception of Time in Music in Patients with Parkinson’s Disease–The Processing of Musical Syntax Compensates for Rhythmic Deficits. Front. Neurosci. 11, 1–11 (2017).

56. Breska, A. & Ivry, R. B. Double dissociation of single-interval and rhythmic temporal prediction in cerebellar degeneration and Parkinson ‘ s disease. PNAS 1–6 (2018) doi:10.1073/pnas.1810596115.

57. Elsinger, C. L. et al. Neural basis for impaired time reproduction in Parkinson’s disease: An fMRI study. J. Int. Neuropsychol. Soc. 9, 1088–1098 (2003).

58. Jones, C. R. G., Malone, T. J. L., Dirnberger, G., Edwards, M. & Jahanshahi, M. Basal ganglia, dopamine and temporal processing: Performance on three timing tasks on and off medication in Parkinson’s disease. Brain Cogn. 68, 30–41 (2008).

59. Merchant, H., Luciana, M., Hooper, C., Majestic, S. & Tuite, P. Interval timing and Parkinson’s disease: Heterogeneity in temporal performance. Exp. Brain Res. 184, 233–248 (2008).

60. O’Boyle, D. J., Freeman, J. S. & Cody, F. W. J. The accuracy and precision of timing of self-paced, repetitive movements in subjects with Parkinson’s disease. Brain 119, 51–70 (1996).

61. Vikene, K., Skeie, G.-O. & Specht, K. Abnormal phasic activity in saliency network, motor areas, and basal ganglia in Parkinson’s disease during rhythm perception. Hum. Brain Mapp. 1–12 (2018) doi:10.1002/hbm.24421.

62. Vikene, K., Skeie, G. O. & Specht, K. Compensatory task-specific hypersensitivity in bilateral planum temporale and right superior temporal gyrus during auditory rhythm and omission processing in Parkinson’s disease. Sci. Rep. 9, 1–9 (2019).

63. Morris, I. B. et al. Music to one’s ears: Familiarity and music engagement in people with Parkinson’s disease. Front. Neurosci. 13, (2019).

64. Martínez-Molina, N., Mas-Herrero, E., Rodríguez-Fornells, A., Zatorre, R. J. & Marco-Pallarés, J. Neural correlates of specific musical anhedonia. Proc. Natl. Acad. Sci. E7337–E7345 (2016) doi:10.1073/PNAS.1611211113.

65. Martinez-Molina, N., Mas-Herrero, E., Rodríguez-Fornells, A., Zatorre, R. J. & Marco-Pallarés, J. White matter microstructure reflects individual differences in music reward sensitivity. J. Neurosci. 39, 5018–5027 (2019).

66. Seger, C. A. et al. Corticostriatal contributions to musical expectancy perception. J. Cogn. Neurosci. 25, 1062–77 (2013).

67. Mas-Herrero, E., Dagher, A. & Zatorre, R. J. Modulating musical reward sensitivity up and down with transcranial magnetic stimulation. Nat. Hum. Behav. 2, 27–32 (2018).

68. Henry, M. J., Herrmann, B., Kunke, D. & Obleser, J. Aging affects the balance of neural entrainment and top-down neural modulation in the listening brain. Nat. Commun. 8, 15801 (2017).

69. Sauvé, S. A., Bolt, E. L. W., Fleming, D. & Zendel, B. R. The impact of aging on neurophysiological entrainment to a metronome. NeuroReport 30, 730–734 (2019).

70. Al Jaja, A., Grahn, J. A., Herrmann, B. & MacDonald, P. A. The effect of aging, Parkinson’s disease, and exogenous dopamine on the neural response associated with auditory regularity processing. Neurobiol. Aging 89, 71–82 (2020).

71. Krampe, R. T., Engbert, R. & Kliegl, R. The effects of expertise and age on rhythm production: Adaptations to timing and sequencing constraints. Brain Cogn. 48, 179–194 (2002).

72. Turgeon, M. & Wing, A. M. Late onset of age-related difference in unpaced tapping with no age-related difference in phase-shift error detection and correction. Psychol. Aging 27, 1152–1163 (2012).

73. Krampe, R. T., Engbert, R. & Kliegl, R. Age-specific problems in rhythmic timing. Psychol. Aging 16, 12–30 (2001).

74. Krampe, R. T., Kliegl, R. & Mayr, U. Timing, sequencing, and executive control in repetitive movement production. J. Exp. Psychol. Hum. Percept. Perform. 31, 379–397 (2005).

75. Chowdhury, R. et al. Dopamine restores reward prediction errors in old age. Nat. Neurosci. 16, 648–653 (2013).

76. Eppinger, B., Schuck, N. W., Nystrom, L. E. & Cohen, J. D. Reduced Striatal responses to reward prediction errors in older compared with younger adults. J. Neurosci. 33, 9905–9912 (2013).

77. Schott, B. H. et al. Ageing and early-stage Parkinson’s disease affect separable neural mechanisms of mesolimbic reward processing. Brain 130, 2412–2424 (2007).

78. Vink, M., Kleerekooper, I., van den Wildenberg, W. P. M. & Kahn, R. S. Impact of aging on frontostriatal reward processing. Hum. Brain Mapp. 36, 2305–2317 (2015).

79. Di Biase, L. et al. Tremor stability index: A new tool for differential diagnosis in tremor syndromes. Brain 140, 1977–1986 (2017).

80. Hoehn, M. M. & Yahr, M. D. Parkinsonism: Onset, progression, and mortality. Neurology 17, 427–442 (1967).

81. Postuma, R. B. et al. MDS clinical diagnostic criteria for Parkinson’s disease. Mov. Disord. 30, 1591–1601 (2015).

82. Cools, R. & D’Esposito, M. Inverted-U-shaped dopamine actions on human working memory and cognitive control. Biol. Psychiatry 69, e113–e125 (2011).

83. Olsen, L. R., Jensen, D. V., Noerholm, V., Martiny, K. & Bech, P. The internal and external validity of the Major Depression Inventory in measuring severity of depressive states. (2003) doi:10.1017/S0033291702006724.

84. Nasreddine, Z. S. et al. The Montreal Cognitive Assessment, MoCA: a brief screening tool for mild cognitive impairment. J Am Geriatr Soc 53, 695–699 (2005).

85. Goetz, C. G. et al. Movement Disorder Society-Sponsored Revision of the Unified Parkinson’s Disease Rating Scale (MDS-UPDRS): Scale Presentation and Clinimetric Testing Results. Mov. Disord. (2008) doi:10.1002/mds.22340.

86. Wallentin, M., Højlund Nielsen, A., Friis-Olivarius, M., Vuust, C. & Vuust, P. The Musical Ear Test, a new reliable test for measuring musical competence. (2010) doi:10.1016/j.lindif.2010.02.004.

87. Fitch, W. T. & Rosenfeld, A. J. Perception and production of syncopated rhythms. Music Percept. Interdiscip. J. 25, 43–58 (2007).

88. Longuet-Higgins, H. C. & Lee, C. S. The Rhythmic Interpretation of Monophonic Music. Music Percept. Interdiscip. J. 1, 424–441 (1984).

89. Lartillot, O., Toiviainen, P. & Eerola, T. A Matlab Toolbox for Music Information Retrieval. Data Anal. Mach. Learn. Appl. 261–268 (2007) doi:10.1007/978-3-540-78246-9_31.

90. Plomp, R. & Levelt, J. M. Tonal consonance and critical bandwidth. J. Acoust. Soc. Am. 38, 548–560 (1965).

91. Sethares, W. A. Tuning, Timbre, Spectrum, Scale. (Springer Science and Business Media, 2004).

92. Matuschek, H., Kliegl, R., Vasishth, S., Baayen, H. & Bates, D. Balancing Type I error and power in linear mixed models. J. Mem. Lang. 94, 305–315 (2017).

93. Stupacher, J. et al. The sweet spot between predictability and surprise: musical groove in brain, body, and social interactions. Front. Psychol. 1–9 (2022) doi:10.3389/fpsyg.2022.906190.

94. Coull, J. T., Cheng, R.-K. & Meck, W. H. Neuroanatomical and neurochemical substrates of timing. Neuropsychopharmacology 36, 3–25 (2011).

95. Matell, M. S. & Meck, W. H. Cortico-striatal circuits and interval timing: coincidence detection of oscillatory processes. Cogn. Brain Res. 21, 139–70 (2004).

96. Teghil, A., Boccia, M., Antonio, D., Vita, A. D. & Lena, C. De. Neural substrates of Internally-Based and Externally-Cued Timing: an activation likelihood estimation (ALE) meta-analysis of fMRI studies. Neurosci. Biobehav. Rev. 96, 197–209 (2019).

97. Cheng (鄭子含), T.-H. Z., Creel, S. C. & Iversen, J. R. How do you feel the rhythm: Dynamic motor-auditory interactions are involved in the imagination of hierarchical timing. J. Neurosci. JN-RM- 1121–21 (2021) doi:10.1523/jneurosci.1121-21.2021.

98. Cope, T. E., Grube, M., Singh, B., Burn, D. J. & Griffiths, T. D. The basal ganglia in perceptual timing: timing performance in Multiple System Atrophy and Huntington’s disease. Neuropsychologia 52, 73–81 (2014).

99. Merchant, H. et al. Finding the beat: a neural perspective across humans and non-human primates. Philos. Trans. R. Soc. 370, 1–16 (2015).

100. Teki, S., Grube, M. & Griffiths, T. D. A unified model of time perception accounts for duration-based and beat-based timing mechanisms. Front. Integr. Neurosci. 5, 90 (2012).

101. Thaut, M. H., Demartin, M. & Sanes, J. N. Brain networks for integrative rhythm formation. PLoS ONE 3, 1–10 (2008).

102. Kung, S.-J., Chen, J. L., Zatorre, R. J. & Penhune, V. B. Interacting cortical and basal ganglia networks underlying finding and tapping to the musical beat. J. Cogn. Neurosci. 25, 401–20 (2013).

103. Morillon, B., Hackett, T. A., Kajikawa, Y. & Schroeder, C. E. Predictive motor control of sensory dynamics in auditory active sensing. Curr. Opin. Neurobiol. 31C, 230–238 (2015).

104. Schubotz, R. I. Prediction of external events with our motor system: towards a new framework. Trends Cogn. Sci. 11, 211–218 (2007).

105. Rimmele, J. M., Morillon, B., Poeppel, D. & Arnal, L. H. Proactive Sensing of Periodic and Aperiodic Auditory Patterns. Trends Cogn. Sci. 22, 870–882 (2018).

106. Arnal, L. H. Predicting “when” using the motor system’s beta-band oscillations. Front. Hum. Neurosci. 6, 1–3 (2012).

107. Nobre, A. C. & van Ede, F. Anticipated moments: temporal structure in attention. Nat. Rev. Neurosci. 19, 34–48 (2018).

108. Oudeyer, P. Y., Gottlieb, J. & Lopes, M. Intrinsic motivation, curiosity, and learning: Theory and applications in educational technologies. Progress in Brain Research vol. 229 (Elsevier B.V., 2016).

109. Schmidhuber, J. Formal Theory of Creativity & Intrinsic Motivation (1990-2010). IEEE Trans. Auton. Ment. Dev. 2, 1–18 (2010).

110. Lumaca, M., Haumann, N. T., Brattico, E., Grube, M. & Vuust, P. Weighting of neural prediction error by rhythmic complexity: a predictive coding account using Mismatch Negativity. Eur. J. Neurosci. 49, 1597–1609 (2019).

111. Vuust, P. et al. To musicians, the message is in the meter: Pre-attentive neuronal responses to incongruent rhythm are left-lateralized in musicians. NeuroImage 24, 560–4 (2005).

112. Drake, C., Penel, A. & Bigand, E. Tapping in time with mechanically and expressively performed music. Music Percept. 18, 1–23 (2000).

113. Palmer, C. & Krumhansl, C. L. Mental Representations for Musical Meter. J. Exp. Psychol. Hum. Percept. Perform. 16, 728–741 (1990).

114. Perna, F., Pavani, F., Zampini, M. & Mazza, V. Behavioral Dynamics of Rhythm and Meter Perception: The Effect of Musical Expertise in Deviance Detection. Timing Time Percept. 6, 32–53 (2018).

115. Geiser, E., Sandmann, P., Jancke, L. & Meyer, M. Refinement of metre perception – training increases hierarchical metre processing. Eur. J. Neurosci. 32, 1979–1985 (2010).

116. Jongsma, M. L. A., Desain, P. & Honing, H. Rhythmic context influences the auditory evoked potentials of musicians and nonmusicians. Biol. Psychol. 66, 129–152 (2004).

117. Vuust, P., Ostergaard, L., Pallesen, K. J., Bailey, C. & Roepstorff, A. Predictive coding of music -Brain responses to rhythmic incongruity. Cortex 45, 80–92 (2009).

118. Hammond, C., Bergman, H. & Brown, P. Pathological synchronization in Parkinson’s disease: networks, models and treatments. Trends Neurosci. 30, 357–364 (2007).

119. Jenkinson, N. & Brown, P. New insights into the relationship between dopamine, beta oscillations and motor function. Trends Neurosci. 34, 611–618 (2011).

120. Fujioka, T., Ross, B. & Trainor, L. J. Beta-Band Oscillations Represent Auditory Beat and Its Metrical Hierarchy in Perception and Imagery. J. Neurosci. 35, 15187–15198 (2015).

121. Fujioka, T., Trainor, L. J., Large, E. W. & Ross, B. Internalized timing of isochronous sounds is represented in neuromagnetic β oscillations. J. Neurosci. 32, 1791–802 (2012).

122. Bauer, A. K. R., Kreutz, G. & Herrmann, C. S. Individual musical tempo preference correlates with EEG beta rhythm. Psychophysiology 52, 600–604 (2015).

123. Iversen, J. R., Repp, B. H. & Patel, A. D. Top-down control of rhythm perception modulates early auditory responses. Ann. N. Y. Acad. Sci. 1169, 58–73 (2009).

124. Graber, E. & Fujioka, T. Endogenous Expectations for Sequence Continuation after Auditory Beat Accelerations and Decelerations Revealed by P3a and Induced Beta-Band Responses. Neuroscience 413, 11–21 (2019).

125. Fujioka, T. & Ross, B. Beta-band oscillations during passive listening to metronome sounds reflect improved timing representation after short-term musical training in healthy older adults. Eur. J. Neurosci. 46, 2339–2354 (2017).

126. Doelling, K. B. & Poeppel, D. Cortical entrainment to music and its modulation by expertise. Proc. Natl. Acad. Sci. 112, E6233–E6242 (2015).

127. te Woerd, E. S., Oostenveld, R., de Lange, F. P. & Praamstra, P. Impaired auditory-to-motor entrainment in Parkinson’s disease. J. Neurophysiol. 117, 1853–1864 (2017).

128. te Woerd, E. S., Oostenveld, R., Lange, F. P. D. & Praamstra, P. Entrainment for attentional selection in Parkinson’s disease. Cortex 99, 166–178 (2018).

129. Gulberti, A. et al. Predictive timing functions of cortical beta oscillations are impaired in Parkinson’s disease and influenced by L-DOPA and deep brain stimulation of the subthalamic nucleus. NeuroImage Clin. 9, 436–449 (2015).

130. Kühn, A. A. et al. Pathological synchronisation in the subthalamic nucleus of patients with Parkinson’s disease relates to both bradykinesia and rigidity. Exp. Neurol. 215, 380–387 (2009).

131. Dauer, W. & Przedborski, S. Parkinson’s Disease: Mechanisms and Models. Neuron 39, 889–909 (2003).

132. Witek, M. A. G. et al. Musicians and non-musicians show different preference profiles for single chords of varying harmonic complexity. Plos One 18, e0281057 (2023).

133. Gothwal, M., Arumugham, S., Yadav, R., Pal, P. & Hegde, S. Deficits in emotion perception and cognition in patients with Parkinson’s disease: A systematic review. Ann. Indian Acad. Neurol. 25, 367 (2022).

134. Lima, C. F., Garrett, C. & Castro, S. L. Not all sounds sound the same: Parkinson’s disease affects differently emotion processing in music and in speech prosody. J. Clin. Exp. Neuropsychol. 35, 373–392 (2013).

135. van Tricht, M. J., Smeding, H. M. M., Speelman, J. D. & Schmand, B. A. Impaired emotion recognition in music in Parkinson’s disease. Brain Cogn. 74, 58–65 (2010).

136. Arnal, L. H. & Giraud, A.-L. Cortical oscillations and sensory predictions. Trends Cogn. Sci. 16, 390–8 (2012).

137. Abbasi, O. & Gross, J. Beta-band oscillations play an essential role in motor–auditory interactions. Hum. Brain Mapp. 1–10 (2019) doi:10.1002/hbm.24830.

