## Supplementary Information for "Dopamine dysregulation in Parkinson’s disease flattens the pleasurable urge to move to musical rhythms" (from another version of this preprint: version 1, 2023-02-27)

**Table S1. ANOVA Results for Wanting to Move Ratings**

| Effect | df | F-value | p-value |
|---|---|---|---|
| Rhythm | 2, 23.5 | 42.16 | < 0.001 |
| Harmony | 2, 25.2 | 96.09 | < 0.001 |
| Group | 3, 94.1 | 0.78 | 0.508 |
| Years of education | 1, 94.0 | 0.51 | 0.477 |
| Years of music training | 1, 94.0 | 0.40 | 0.527 |
| MDI | 1, 94.0 | 0.80 | 0.374 |
| Rhythm:Harmony | 4, 4909.6 | 3.57 | 0.007 |
| Rhythm:Group | 6, 97.0 | 21.22 | < 0.001 |
| Harmony:Group | 6, 97.0 | 18.37 | < 0.001 |
| Rhythm:Harmony:Group | 12, 4909.6 | 1.65 | 0.072 |

**Table S2. Effects of Rhythmic Complexity on Wanting to Move Ratings**

| Group | Contrast | Estimate | df | 95% CI |
|---|---|---|---|---|
| $HC_{young}$ | Linear | -1.469 | 80.0 | -1.849, -1.089 |
| $HC_{young}$ | Quadratic | -2.556 | 77.5 | -3.202, -1.909 |
| $HC_{old}$ | Linear | -0.512 | 80.0 | -0.892, -0.133 |
| $HC_{old}$ | Quadratic | -1.340 | 77.5 | -1.986, -0.693 |
| $PD_{DA}$ | Linear | -0.126 | 87.1 | -0.528, 0.277 |
| $PD_{DA}$ | Quadratic | -0.227 | 84.9 | -0.911, 0.456 |
| $PD_{nonDA}$ | Linear | -0.468 | 85.3 | -0.864, -0.072 |
| $PD_{nonDA}$ | Quadratic | -0.273 | 83.0 | -0.946, 0.400 |

**Table S3. Effects of Harmonic Complexity on Wanting to Move Ratings**

| Group | Contrast | Estimate | df | 95% CI |
|---|---|---|---|---|
| $HC_{young}$ | Linear | -0.037 | 92.8 | -0.325, 0.251 |
| $HC_{young}$ | Quadratic | -0.407 | 82.3 | -0.814, -0.001 |
| $HC_{old}$ | Linear | -0.895 | 92.8 | -1.183, -0.607 |
| $HC_{old}$ | Quadratic | -1.451 | 82.3 | -1.857, -1.044 |
| $PD_{DA}$ | Linear | -0.829 | 94.9 | -1.140, -0.517 |
| $PD_{DA}$ | Quadratic | -0.162 | 87.0 | -0.599, 0.275 |
| $PD_{nonDA}$ | Linear | -0.995 | 94.4 | -1.300, -0.690 |
| $PD_{nonDA}$ | Quadratic | -0.190 | 85.9 | -0.618, 0.239 |

**Table S4. ANOVA Results for Pleasure Ratings**

| Effect | df | F-value | p-value |
|---|---|---|---|
| Rhythm | 2, 30.2 | 19.88 | < 0.001 |
| Harmony | 2, 28.5 | 52.14 | < 0.001 |
| Group | 3, 94.0 | 3.49 | 0.019 |
| Years of education | 1, 94.0 | 2.94 | 0.090 |
| Years of music training | 1, 94.0 | 0.68 | 0.411 |
| MDI | 1, 94.0 | 0.019 | 0.890 |
| Rhythm:Harmony | 4, 4896.8 | 2.56 | 0.037 |
| Rhythm:Group | 6, 119.9 | 6.79 | < 0.001 |
| Harmony:Group | 6, 98.8 | 7.60 | < 0.001 |
| Rhythm:Harmony:Group | 12, 4896.8 | 2.06 | 0.016 |

**Table S5. Effects of Rhythmic Complexity on Pleasure Ratings**

| Group | Contrast | Estimate | df | 95% CI |
|---|---|---|---|---|
| $HC_{young}$ | Linear | -1.243 | 112.3 | -1.697, -0.788 |
| $HC_{young}$ | Quadratic | -1.778 | 87.9 | -2.525, -1.031 |
| $HC_{old}$ | Linear | -0.375 | 112.3 | -0.829, 0.080 |
| $HC_{old}$ | Quadratic | -0.650 | 87.9 | -1.397, 0.097 |
| $PD_{DA}$ | Linear | -0.097 | 122.1 | -0.580, 0.387 |
| $PD_{DA}$ | Quadratic | -0.246 | 94.5 | -1.039, 0.547 |
| $PD_{nonDA}$ | Linear | -0.602 | 119.6 | -1.077, -0.127 |
| $PD_{nonDA}$ | Quadratic | -0.528 | 92.8 | -1.308, 0.252 |

**Table S6. Effects of Harmonic Complexity on Pleasure Ratings**

| Group | Contrast | Estimate | df | 95% CI |
|---|---|---|---|---|
| $HC_{young}$ | Linear | -0.021 | 103.3 | -0.365, 0.324 |
| $HC_{young}$ | Quadratic | -0.062 | 79.5 | -0.440, 0.316 |
| $HC_{old}$ | Linear | -0.951 | 103.3 | -1.295, -0.606 |
| $HC_{old}$ | Quadratic | -0.502 | 79.5 | -0.880, -0.124 |
| $PD_{DA}$ | Linear | -0.845 | 104.2 | -1.218, -0.473 |
| $PD_{DA}$ | Quadratic | -0.174 | 86.0 | -0.580, 0.232 |
| $PD_{nonDA}$ | Linear | -0.935 | 104.0 | -1.300, -0.571 |
| $PD_{nonDA}$ | Quadratic | -0.083 | 84.4 | -0.481, 0.315 |
